# Heat stress at the bicellular stage inhibits sperm cell development and their transport into pollen tubes

**DOI:** 10.1101/2023.09.13.557624

**Authors:** Xingli Li, Astrid Bruckmann, Thomas Dresselhaus, Kevin Begcy

## Abstract

For a successful double fertilization process in flowering plants (angiosperms), pollen tubes each deliver two non-motile sperm cells towards female gametes (egg and central cell, respectively). Heatwaves especially during the reproduction period are threatening male gametophyte (pollen) development, which results in severe yield losses. By using maize as a crop and grass model system, we found strong seed set reduction when moderate heat stress was applied for two days during the uni- and bicellular stages of pollen development. We show that heat stress accelerates pollen development and impairs pollen germination capabilities, when applied at the unicellular stage. Heat stress at the bicellular stage impairs sperm cell development and their transport into pollen tubes. To understand the course of the latter defects, we used marker lines and analyzed the transcriptomes of isolated sperm cells. While heat stress also affects the expression of genes involved in transcription, RNA processing and translation, especially genes in DNA replication and the cell cycle were mis-regulated. This includes centromeric histone CENH3 and α-tubulin. Most mis-regulated genes are involved in transition from metaphase to anaphase during pollen mitosis II (PM II). Heat stress activates spindle assembly check point and meta-to anaphase transition genes in sperm cells. In summary, mis-regulation of the identified genes during heat stress at the bicellular stage explains sperm cell development and transport defects ultimately leading to sterility.

## INTRODUCTION

Global warming is associated with hotter, longer, and more frequent heat waves. When these high temperature episodes occur during reproductive development in crop plants, a significant decrease in yield is commonly observed (Folsom et al., 2014; Chen et al., 2016; Begcy et al., 2019). Within reproductive development, male gametophyte formation is one of the most susceptible stages (De Storme and Geelen, 2014; Begcy et al., 2019; Chaturvedi et al., 2021). In flowering plants (angiosperms), male gametophyte development undergoes two phases, microsporogenesis and microgametogenesis. Four microspores are generated during the first phase from a microspore mother cell undergoing meiosis. During the second phase, two rounds of mitotic division give rise to tricellular pollen grains in most angiosperm (Zhou et al., 2017; Hafidh and Honys, 2021; Huang et al., 2021). The first division, known as pollen mitosis I (PMI), is highly asymmetric and produces a small generative cell and a large vegetative cell. During pollen mitosis II (PMII) the small generative cell goes through a symmetric division forming two sperm cells engulfed by the large vegetative cell that enters into the G0 phase of the cell cycle. Thus, a mature pollen grain in most angiosperms contains two small sperm cells attached to the nucleus of the large vegetative cell (Borg et al., 2009; Long et al., 2019; Huang et al., 2021).

In angiosperms including maize, sperm cells have lost their motility and are transported as a passive cargo through the pollen tube towards the ovule for double fertilization (Zhou and Dresselhaus, 2019). After pollen germination, sperm cells move as a male germ unit (MGU) attached to the nucleus of the vegetative cell before being delivered in the receptive synergid cell and separated (Dumas et al., 1985; Dresselhaus et al., 2016; Sprunck, 2020; Sugi et al., 2023). Sperm cell transport and delivery requires a well-organized and dynamic cytoskeleton system (Cheung et al., 2008; Sharma et al., 2021). Using fluorescent labeled actin-binding protein and other pollen specific markers have been demonstrated that pollen tube growth is especially dependent on the dynamic organization and regulation of actin and its microfilaments (Cheung et al., 2008; Chang et al., 2013; Takeuchi and Higashiyama, 2016; Zhang et al., 2023). In tobacco and *Arabidopsis* pollen tubes, it has been shown that heat stress alters cytoskeletal dynamics, isoform content, and spindle orientation (De Storme and Geelen, 2013; Parrotta et al., 2016).

The symmetric division occurring during PMII is fundamental for the generation of the two sperm cells that will arise from the generative cell either within the pollen grain in most angiosperms or after germination inside the pollen tube like in tobacco and *Amborella trichopoda* (Durbarry et al., 2005; Flores-Tornero et al., 2020; Huang et al., 2021). In *Arabidopsis*, it was shown that sperm cells are not necessary and do not control pollen tube growth and guidance (Zhang et al., 2017).

To ensure accurate chromosome segregation during germ cell division and to produce two genetically identical cells during PM II, while further cell division of the vegetative cell is prevented, highly accurate cell cycle checkpoint mechanisms must be executed along these processes. During mitosis, the spindle-assembly checkpoint (SAC) is one of these mechanisms that safeguards the transition from metaphase to anaphase (Lara-Gonzalez et al., 2021; Farrell et al., 2023). SAC monitors proper attachment of spindle microtubules to the surface of the kinetochores. In *Arabidopsis*, SAC architecture is different from the one characterized in yeast and animals (Caillaud et al., 2009; Komaki and Schnittger, 2016; Komaki and Schnittger, 2017). Moreover, plant cells are able to reset the cell cycle with duplicated chromosomes avoiding nuclear division (Kozgunova et al., 2019). In maize, some of the major components of the SAC have been identified and showed conserved function (Yu et al., 1999; Du and Dawe, 2007; Li and Dawe, 2009; Su et al., 2021), but it is unclear whether SAC plays also a role in PM II. Reduction in fertilization success and yield due to defects in male gametophyte performance have been largely attributed to the lack of pollen viability or inability to germinate under high temperatures (Firon et al., 2006; Endo et al., 2009; De Storme and Geelen, 2014; Wang et al., 2017; Begcy et al., 2018; Begcy et al., 2019). Low starch content, decreased enzymatic activity, energy, and lipid formation are major cellular components impacted by temperature (Thalmann and Santelia, 2017; Yang et al., 2018; Begcy et al., 2019; Kumar et al., 2023). In a previous study, we have shown that transient heat stress over two days at the tetrad stage has a strong impact on further pollen performance (Begcy et al., 2019). It remained unclear whether successive stages, uni- and bicellular pollen, as well as sperm cell formation and delivery are also affected if heat stress is applied at later stages.

Here, we demonstrate that transient heat stress during the uni- and bicellular stages accelerates pollen development in maize. Moreover, sperm cell formation and their transport in the pollen tube is affected ultimately leading to sterility. With a focus on heat stress application during the bicellular stage, we aimed to understand the underlying molecular mechanisms that lead to sterility after comparing the transcriptomes and proteomes of stressed and un-stressed pollen grains.

## RESULTS

### Seed set is strongly reduced if heat stress is applied during the uni- or bicellular stage of pollen development

Historically, pollen development studies under heat stress are performed across several developmental stages simultaneously and occasionally through the entire pollen formation process (Firon et al., 2006; Wang et al., 2017; Begcy et al., 2019; Wada et al., 2020). By dissecting the impact of heat stress on a single pollen stage, we previously showed that heat stress during the tetrad stage impacts starch, lipid, and energy metabolism (Begcy et al., 2019). To elucidate whether heat stress triggers similar responses during other stages of pollen development, we imposed heat stress (35°C/25°C light/dark period) for 48 h on maize plants, specifically either at the unicellular and the bicellular stage, respectively. A parallel set of maize plants was maintained under optimal growth conditions (25°C/21°C light/dark period) in an equivalent chamber and was used as a control for all experiments (see Fig. 1A for experimental setup). Since fertilization ability is the most important indicator to evaluate pollen quality, we first pollinated non-stressed (NS) cobs with NS pollen as a control and as expected obtained full seed set (Fig. 1B-C). Contrary, cobs pollinated with pollen heat stressed (HS) only at the unicellular stage (Fig. 1B) and at the bicellular stage (Fig. 1C), respectively, resulted in severe reduction of seed set (Fig. 1B-C). Taken together these results show that heat stress applied during either the unicellular or the bicellular stage of pollen development leads to sterility and strongly reduced yield.

**Figure 1.**
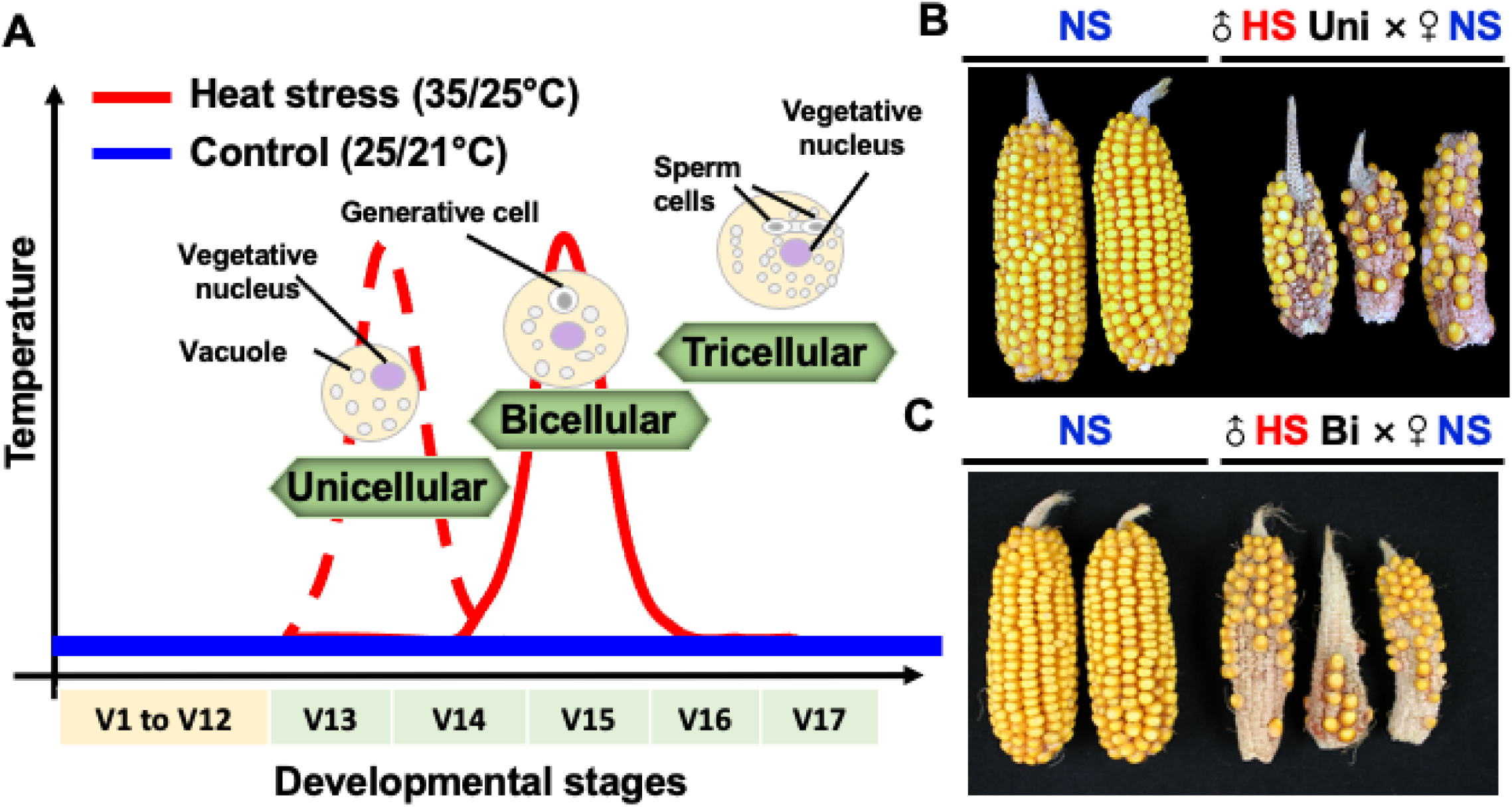
Heat stress during the unicellular and bicellular stages of pollen development reduces seed set in maize. **(A)** Experimental setup. Maize plants were grown in control conditions (25°C/21°C light/dark period) until they reached either the unicellular and bicellular stage of pollen development. Stages were identified according to Begcy and Dresselhaus (2017). At stage V13 (unicellular; red dotted line) and V15 (bicellular; red line), respectively, plants were exposed to moderate heat stress (35°C/25°C light/dark period) for 48 h and afterward transferred back to control conditions until maturity. Blue line indicates (non-stressed) conditions. Non-stressed cobs were pollinated with both, non-stressed pollen (NS × NS) and **(B)** heat-stressed pollen at the unicellular stage (♂HS Uni × ♀NS) and **(C)** heat-stressed pollen at the bicellular stage (♂HS Bi × ♀NS), respectively. Seed set was strongly reduced after pollinating with heat-stressed pollen at both pollen developmental stages.

### Heat stress at the uni- and bicellular stages accelerates pollen development

To further explored the possible causes of seed set reduction, we initially analyzed the effect of HS on pollen morphology and viability immediately after HS application. Similar as observed during HS application at the tetrad stage, we found decreased pollen viability when HS was applied at the unicellular (Supplemental Fig. 1), but not at the bicellular stage (Supplemental Fig. 2). However, in contrast to HS during the tetrad stage, we did not see any morphological alteration after HS application at both stages (Supplemental Figs. 1 and 2). Fluorescein Diacetate (FDA) was used as a proxy to measure pollen enzymatic activity of pollen stressed at the unicellular stage. In contrast to NS pollen (Fig. 2A), HS pollen showed significantly lower enzymatic activity (Fig. 2B and C). To further explore the impact of heat stress on the timing of pollen development, we stained NS (Fig. 2D) and HS pollen (Fig. 2E) at the unicellular stage with DAPI (4′,6-diamidino-2-phenylindole). More than half of the NS pollen (63%) were at the unicellular stage, whereas only 30% of HS pollen were at the unicellular stage (Fig. 2F). A large portion of them progressed to the bicellular (35%) and tricellular (11%) stages (Fig. 2E and F), respectively.

**Figure 2.**
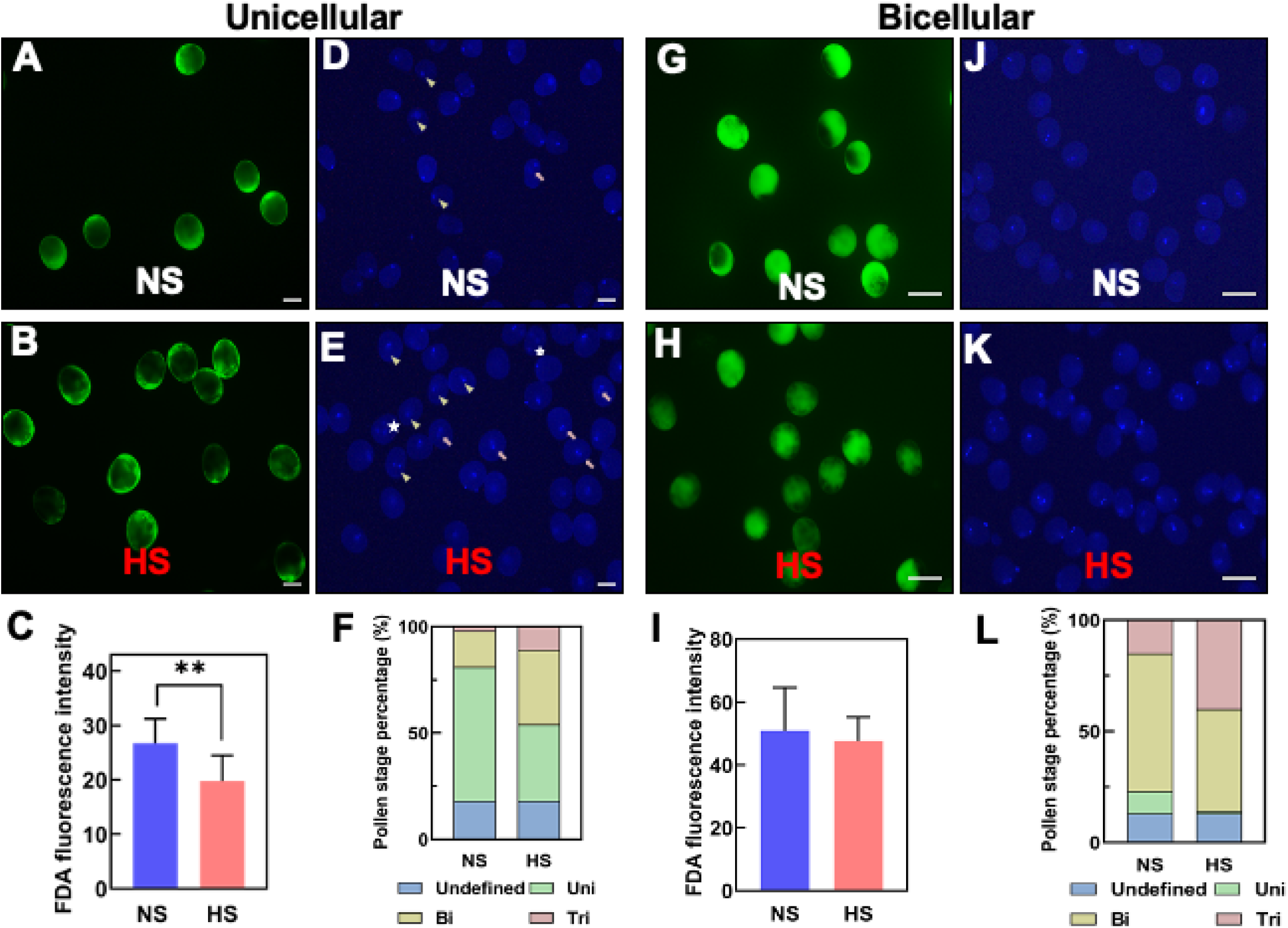
Heat stress at the unicellular and bicellular stages accelerates pollen development. (A) Pollen stained with FDA after non-stressed (NS) and **(B)** heat-stressed (HS) treatment at the unicellular stage. **(C)** Quantification of enzymatic activity of pollen shown in (A and B). **(D)** DAPI staining of NS and **(E)** HS pollen as in (A and B). Triangles: unicellular stage, arrows: bicellular stage and stars: tricellular stage. **(F)** Quantification of pollen shown in (D and E). Blue: undefined stage, green: unicellular stage, yellow: bicellular stage and red: tricellular stage of maize pollen development. **(G)** Pollen stained with FDA after non-stressed (NS) and heat-stressed (HS) treatment at the bicellular stage. **(I)** Quantification of enzymatic activity of pollen shown in (G and H). **(J)** DAPI staining of NS and **(K)** HS pollen as in (G and H). **(L)** Quantification of pollen shown in (J and K). For description see (F). Data are presented as the mean ± SD. n = 400–500. Scale bars = 50 μm. Two asterisks indicate significant difference at P < 0.001; one-tailed *t*-test comparing heat-stressed (HS - red) to non-stressed (NS - blue) samples.

When we applied HS at the bicellular stage of pollen development, we could not see any significant differences between NS and HS pollen after FDA staining (Fig. 2G-I). After DAPI staining (Fig. 2J and K), we observed that pollen developed in NS conditions were mostly (65%) at the bicellular stage. Pollen at the unicellular (10%) and tricellular (15%) stage were also observed (Fig. 2L). In contrast, HS pollen showed reduced pollen numbers at uni-cellular (1%) and bicellular stages (46%). Many pollen were already at the tricellular (39%) stage (Fig. 2L). These results indicate that heat stress accelerates the timing of pollen development and has only a minor effect on pollen vitality and only when applied at the unicellular stage.

### Heat stress at the unicellular stage impairs pollen germination capabilities

To further explore the impact of HS on pollen development during both stages, we transferred HS plants to NS conditions and allowed plants to recover and continue pollen development until maturation. Using mature pollen, we studied *in vitro* and *in vivo* germination and pollen tube growth capabilities of both, NS and HS pollen at the unicellular (Fig. 3A, B, E and F) and bicellular stage (Fig. 3G, H, J and K). Quantification of *in vitro* germination rate (Fig. 3C) and speed (Fig. 3D) after HS application at the unicellular stage showed both, a reduction of germination rate of approximately 83% to around 63%, as well as a significant reduction in pollen tube growth speed. To further confirm these findings, we conducted *in vivo* germination experiments to investigate the pollen penetration ability on maize silks (Fig. 3E and F). While NS pollen germinated on silks and penetrated the transmitting track (Fig. 3E), HS pollen at the unicellular stage attached to the silk but were unable to grow a pollen tube into the silk (Fig. 3F).

**Figure 3.**
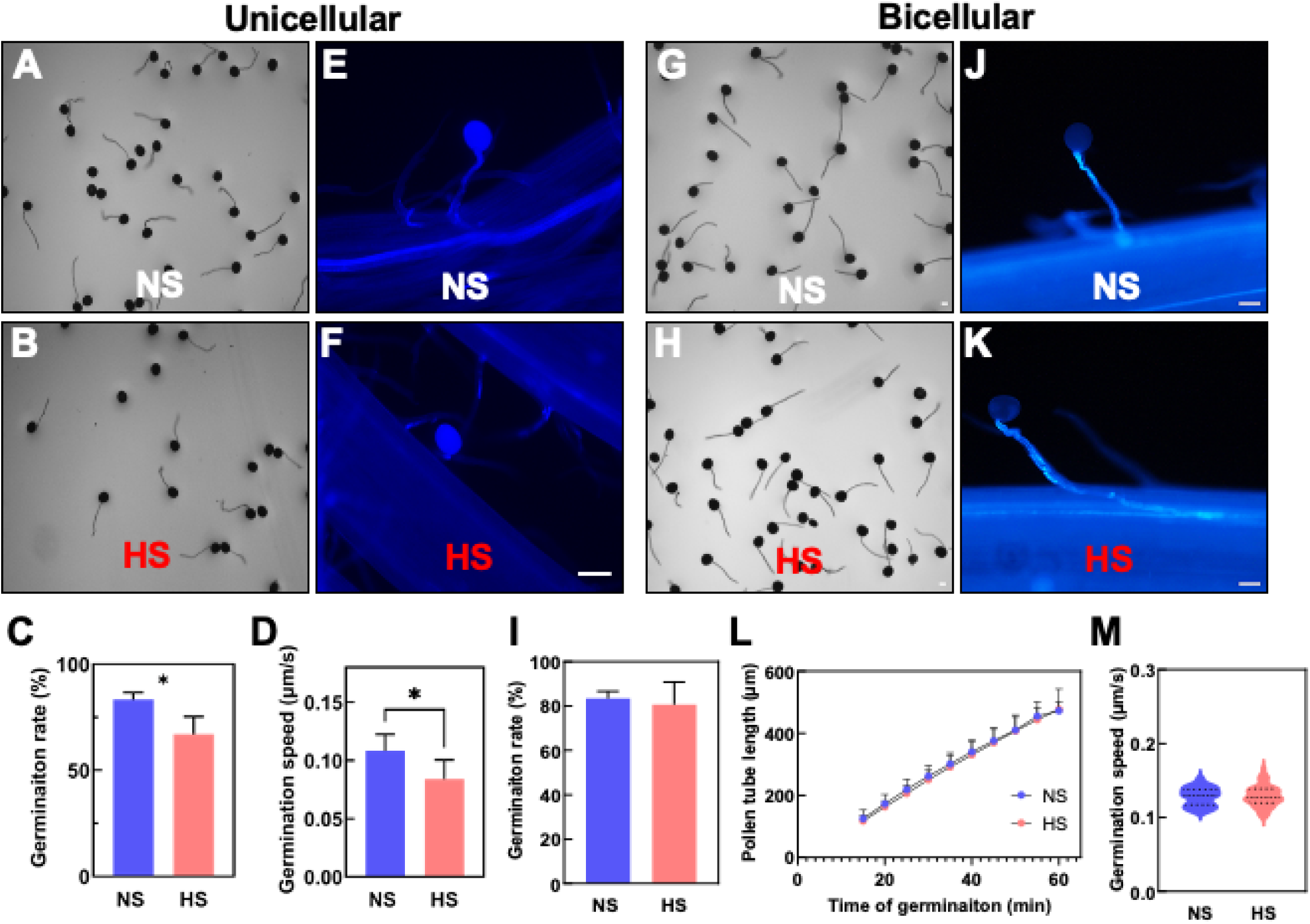
Heat stress applied at the unicellular, but not at the bicellular stage impairs pollen germination capabilities. **(A)** *In vitro* germination assays of pollen isolated from NS and **(B)** HS plants stressed at the unicellular stage. (C) *In vitro* germination rate of pollen shown in (A and B). **(D)** Germination speed of NS and HS pollen stressed at the unicellular stage. **(E)** Aniline blue staining of NS and **(F)** HS pollen stressed at the unicellular stage germinating on papilla hair cells. (G) *In vitro* germination assays of pollen from NS and **(H)** HS plants stressed at the bicellular stage. (I) *In vitro* germination rate of pollen harvested from NS and HS plants stressed at the bicellular stage show no germination rate differences. **(J)** Aniline blue staining of NS and **(K)** HS pollen stressed at the bicellular stage germinating on papilla hair cells show normal pollen tube penetration. **(L)** Pollen tube length and **(M)** germination speed of NS and HS pollen stressed at the bicellular stage. Statically significant differences in germination rate, pollen tube length and germination speed could not be detected between both conditions. Scale bars = 100 μm. Data are presented as the mean ± SD. n = 400–500. One asterisk indicates significant difference at P < 0.01; one-tailed *t*-test comparing HS to NS samples.

We did not observe such differences in pollen germination and growth capabilities when HS was applied at the bicellular stage. Pollen germination rate *in vitro, in vivo,* silk penetration, and growth was similar to the NS control (Fig. 3G-M). In conclusion, the observed sterility (Fig. 1B) when HS was applied at the unicellular stage could be explained by pollen germination, growth, and penetration defects. However, sterility caused after HS exposure during the bicellular stage remained unclear.

### Heat stress at the bicellular stage impairs sperm cell development and transport

To further elucidate the causes of the strong seed set reduction phenotype without noticeable changes in morphological, cellular, and biochemical properties of HS pollen at the bicellular stage, we next explored the developmental processes occurring at the transition between the bicellular to the tricellular stage of maize pollen development. During the transition to the tricellular stage, the generative cell undergoes mitosis giving rise to two sperm cells which during fertilization will fuse with the central cell and the egg cell to form the endosperm and the embryo, respectively (Zhao et al., 2017). Therefore, since the formation of the two sperm cells is the main and critical developmental process occurring at the bicellular stage, we hypothesized that heat stress might impact sperm cell development. To test this hypothesis, we used a maize germ and sperm cell marker line that expresses an α-tubulin gene fused with the yellow florescence protein (YFP) and thus labels the spindle apparatus during germ cell division (Kliwer, et al., 2009). Under NS conditions, the α-tubulin-YFP signal was observed during the mitotic division of the generative cell and marks the two spindle-formed sperm cells after division (Fig. 4A-C). In contrast, the α-tubulin-YFP signal in HS pollen was significantly reduced and the spindle apparatus and sperm cell shape appeared less symmetric (Fig. 4D-F). Quantification of α-tubulin-YFP signals cells confirmed this observation and were about 30% reduced in HS sperm cells compared with NS sperm cells (Fig. 4G). These results demonstrated that even though the morphology of HS pollen did not change, sperm cell formation was significantly impaired.

**Figure 4.**
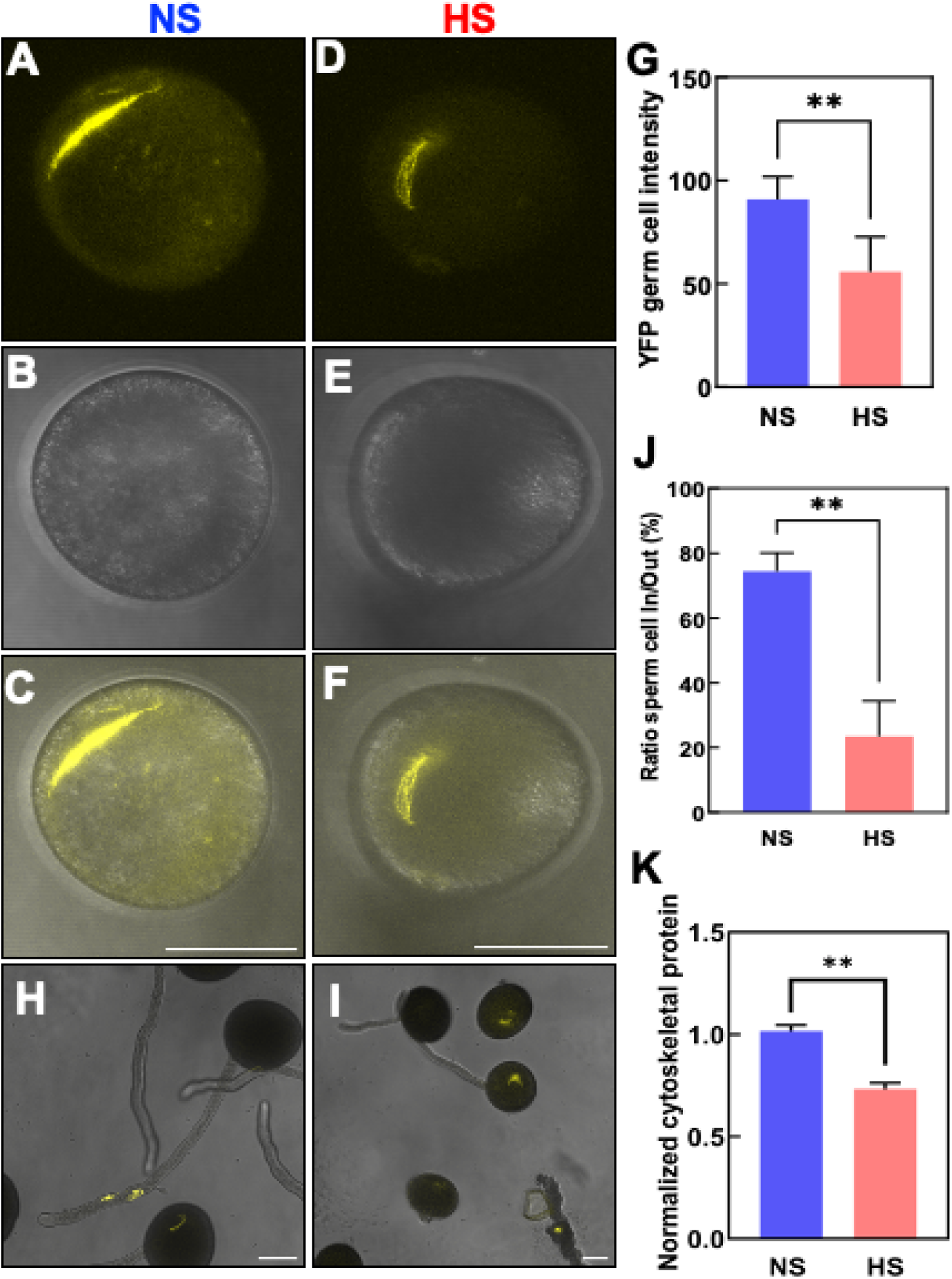
Heat stress at the bicellular stage impairs sperm cell development and their transport into the pollen tube. Maize sperm cell maker lines containing α-tubulin fused with YFP were grown in control conditions (25°C/21°C light/dark period) until they reached the bicellular stage of pollen development. Then plants were exposed to heat stress (35°C/25°C light/dark period) for 48 h. A parallel set of plants was maintained under control conditions. **(A-F)** Confocal images of NS (A-C) and HS pollen (D-F) stressed at the bicellular stage. (C and F) Merged images of (A and B) and (D and E), respectively. **(G)** Quantification of YFP signal intensity of pollen as indicated. **(H)** Confocal images of NS pollen grains showing sperm cells travelling into the pollen tube. **(I)** Sperm cells were kept inside the pollen grain after HS was applied at the bicellular stage of pollen development. **(J)** Ratio of sperm cells remaining and travelling inside the pollen tube as indicated. **(K)** Reduction in the number of detectable cytoskeletal-related proteins in NS and HS pollen stressed at the bicellular stage. See Suppl. Table S1 for details. Asterisks indicate a significant difference at *P* < 0.001; one-tailed *t-*test comparing NS and HS samples. Scale bars = 50 μm. n = 400–500.

We next investigated whether this also impacts sperm cell transport inside pollen tubes. We monitored the sperm cell journey using the α-tubulin-YFP marker line and found that within 1h after pollen germination and growth *in vitro*, about 80% NS sperm cells were visible inside pollen tubes (Fig. 4H and J). In contrast, only about 20% sperm cells were visible inside pollen tubes formed by HS pollen (Fig. 4I and J).

To further verify that the expression of not only one single a-tubulin gene was reduced after heat stress, we performed liquid chromatography with tandem mass spectrometry (LC–MS/MS) from mature pollen (Supplemental Table S1) to gain further insights into the effects of heat stress at the proteome level on cytoskeletal proteins. Only proteins that were detected in all three biological replicates (‘max count 3’) and met the valid threshold of Mascot score >10, peptides >2 in at least one of the conditions (NS and HS) were considered for further analysis. We normalized the number of identified tubulin and actin peptides as well as their associated proteins and noted a significant reduction of tubulin and actin related genes at HS conditions (Fig. 4K). Taken together our cellular and proteomic data show that heat stress at the bicellular stage impairs sperm cell development and transport due to reduced amounts of cytoskeletal tubulin and actin proteins involved in both, sperm cell formation and transport.

### Heat stress at the bicellular stage decreases centromeric histone levels

It remained unclear whether reduced tubulin levels and a malformed spindle apparatus leads to chromosome losses and thus sperm cells defects. The centromeric histone H3 variant CENH3 is essential for kinetochore assembly and establishment, and thus an ideal marker to label each chromosome showing also equivalent homologous chromosome segregation in eukaryotes (Collins et al., 2005; Keçeli et al., 2020). We generated a sperm cell-specific mRuby3-CENH3 marker by using 1.3 kbp upstream of the sperm cell-specific *GEX3* gene of maize (Chen et al., 2017) as promoter. Ten chromosomes were clearly visible in each sperm cell inside NS pollen grains at maturity due to mRuby3-CENH3 labelling of centromeric chromosome regions (Fig. 5A). However, when the GEX3p:mRuby3-CENH3 marker line was exposed to HS conditions during the bicellular stage of pollen development fluorescent signals at some chromosomes appeared weaker, although a comparable number of chromosomes could be observed (Fig. 5B). Quantification of mRuby3-CENH3 signal showed a significant reduction in HS pollen compared to NS conditions (Fig. 5C). These results indicate that CENH3 levels are reduced after heat stress and appear more irregular at centromeric regions indicating sperm cell defects.

**Figure 5.**
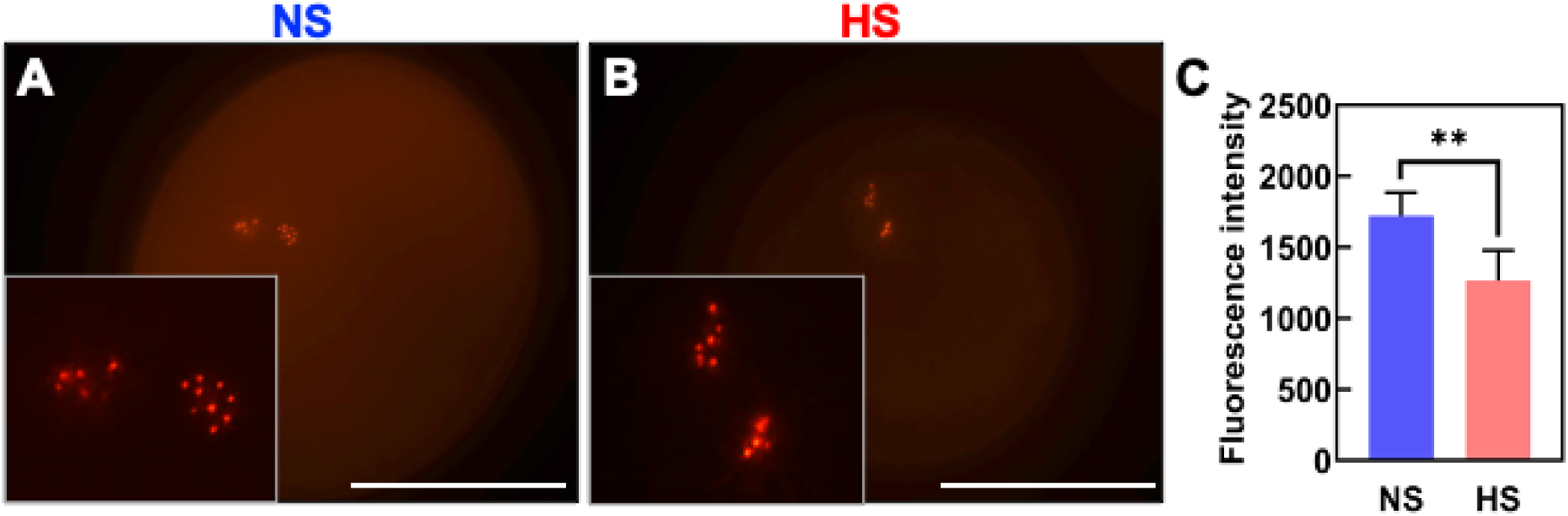
Heat stress during the bicellular stage decreases content of centromeric histones in maize. Marker lines expressing centromeric histones (CENH3) specifically in sperm cells (GEX3p:mRuby3-CENH3) were grown in control conditions (25°C/21°C light/dark period) until they reached the bicellular stage of pollen development and then submitted to heat stress (35°C/25°C light/dark period) for 48 h. A parallel set of marker line plants was maintained under optimal conditions and used as controls. **(A)** Confocal images showing CENH3 in mature pollen grains from NS and **(B)** HS plants. Insets show enlarged sperm cells containing CENH3 signals. **(C)** Intensity quantification of mRuby3-CENH3 in both conditions. Asterisks indicate a significant difference at *P* < 0.001; one-tailed *t-*test comparing HS to NS samples. Scale bars = 50 μm.

### Heat stress impacts expression of genes involved in transcription, DNA replication, RNA processing, and translation in sperm cells

To elucidate the molecular mechanisms impacted by HS during maize sperm cell development, we used an RNA-seq approach to further analyze the transcriptional changes imposed by increased temperatures at the bicellular stage. Maize sperm cells were isolated using a Percoll gradient strategy (Dupuis et al., 1987; Chen et al., 2017). We isolated approximately 5000 individual sperm cells in each of the three biological replicates of both, NS and HS samples (Fig. 6A-B; Supplemental Table S2). We found 952 genes differentially expressed between NS and HS conditions (Fig. 6C). Among these genes, 267 were downregulated and 684 were upregulated (Supplemental Table S3). Gene ontology analysis showed enrichment of genes involved in chromosome and chromatin organization, DNA conformation, cellular homeostasis, and purine-containing compound metabolic process (Supplemental Fig. S3). Previously, we showed that HS at the tetrad stage of maize pollen development induced the expression of several heat shock protein genes (HSPs), which was still detectable in pollen at maturity (Begcy et al., 2019). Thus, we compared expression of HSP genes in sperm cells at both conditions. An overall low or lack of expression of HSP and heat shock factor (HSF) genes have been reported in NS sperm cells of maize (Chen et al., 2017). After HS application, we found a significant transcriptional increase only in HSP11 (Zm00001eb011930), HSP70-3 (Zm00001eb012510) and HSP70-4 (Zm00001eb136490) (Fig. 6D). HSP70-6 (Zm00001eb012470) was not induced in sperm cells after heat stress at the bicellular stage and was used as a control (Fig. 6D). These results suggest that short spikes of heat stress likely do not interfere significantly with the gene regulatory network of sperm cells.

**Figure 6.**
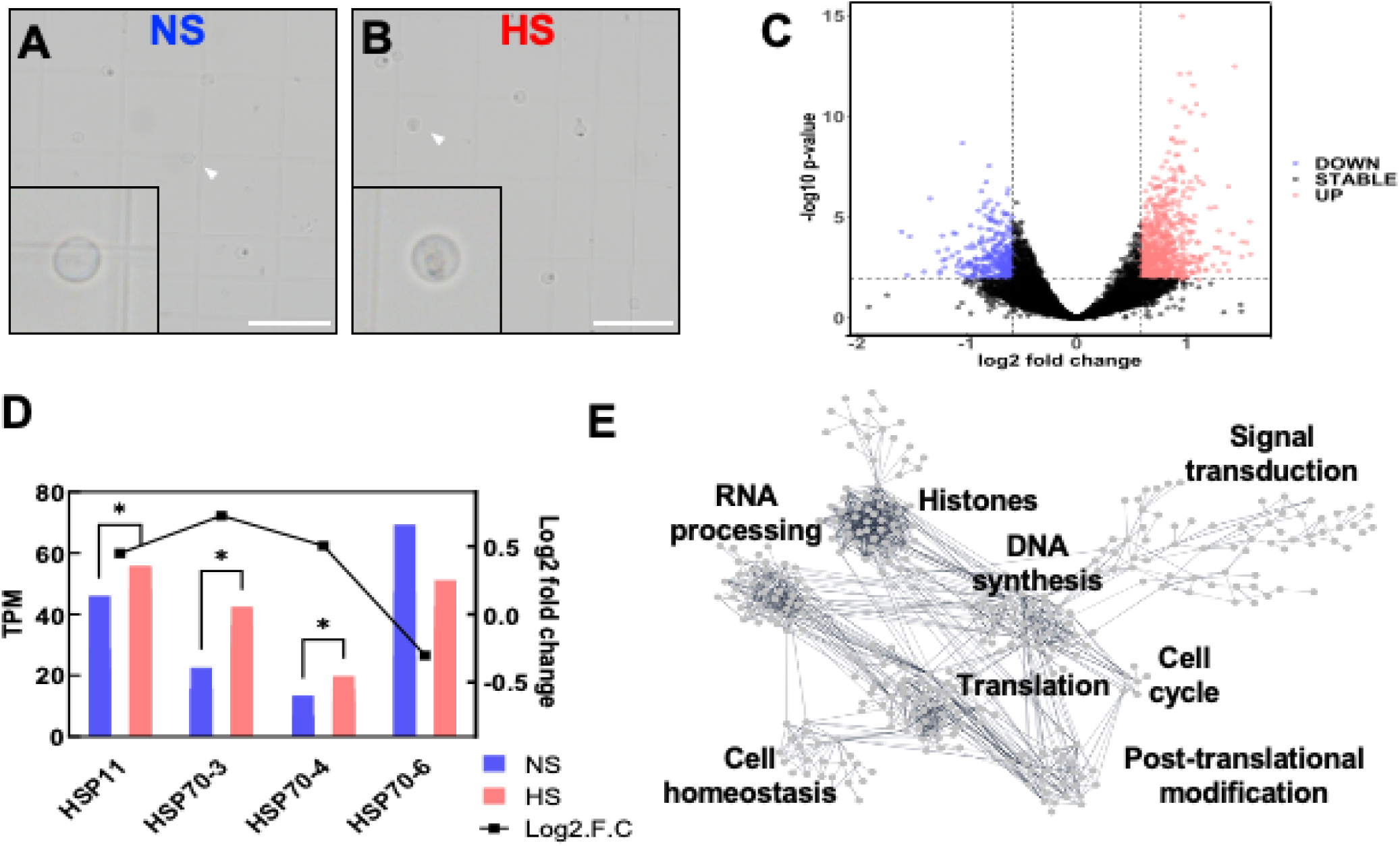
Heat stress at the bicellular stage mis-regulates replication-associated genes in sperm cells. **(A)** Confocal images of NS and **(B)** HS sperm cells stressed at the bicellular stage of pollen development. Insets show enlargement of single sperm cells (arrowheads). Scale bar = 50 μm. Approximately 5,000 individual sperm cells in each of the three biological replicates per conditioned were used. **(C)** Differential gene expression (base 2 logarithm fold change) in sperm cells harvested at maturity after heat stress was applied at the bicellular stage of pollen development. HS samples relative to control (NS) samples were plotted versus average gene expression levels (i.e. logarithm of mean counts normalized for difference in library sizes). Red color indicates upregulation. Blue color indicates downregulation. Black color indicates no significant transcriptional change. **(D)** TPM (transcript per million) vales and RT-qPCR analysis shows that differential expression of HSP genes in sperm cells was still increased in mature pollen. Asterisks indicate significant difference at P < 0.01; one-tailed *t*-test comparing HS to NS samples. n = 3 biological replicates, each with 3 technical replicates. **(E)** Gene network analysis of interactions of differentially expressed genes in sperm cells in response to heat stress at the bicellular stage. A threshold of 0.7 of edge confidence was used. A detailed list of the genes included in the gene interaction analysis can be found in Suppl. Table S4.

To further understand sperm cell development-associated genetic responses to HS, we used the identified differentially expressed genes and conducted a cluster correlation analysis to elucidate functional groups of genes with shared expression profiles. We merged our cluster correlation analysis with the web-tool STRING (Szklarczyk et al., 2019), to identify previously described genes as member of any given pathway. First, we remove unrooted genes and identified 331 interacting genes associated with the HS response in sperm cells (Supplemental Table S4). To select candidate genes in response to HS from sperm cells, we used a stringent edge confidence of at least 0.7. Previously using this approach, we found five main clusters after heat stress during the tetrad stage of pollen development (Begcy et al., 2019). Our current analysis yielded eight main hubs affected by HS in sperm cells (Fig. 6E). Notably, these hubs are formed by genes related to histones, DNA synthesis, RNA processing, translation, post-translational modification, cell cycle and signal transduction (Fig. 6E). Notably, the mis-regulation of histone genes confirms the previous observation that levels of centromeric histone CENH3 is reduced after HS (Fig. 5).

### Heat stress affects highly expressed genes in sperm cells

To compare the transcriptional status of NS and HS sperm cells, we analyzed the genome wide transcript per million (TPM) values of both conditions (Supplemental Fig. S4). When TPM values from the entire maize genomes were plotted, a high correlation (R² = 0.9868) in gene expression was obtained between both conditions (Supplemental Fig. S4A-B), suggesting that even though HS at the bicellular stage impacts the expression of genes in sperm cells, but the level of changes appeared minor. However, this analysis also included a larger portion of genes that are not expressed in any of the conditions. Our sperm cell analysis yielded an average of 30% transcriptional expression of the entire maize genome in both conditions. No significant differences were observed when genes >1 and <100 TPM were used (Supplemental Fig. S4B). However, when we used a 100 TPM cutoff to explore the transcriptional impact of HS on sperm cells, a lower correlation of TPM levels (R² = 0.7477) was found when only truly expressed genes were used in the analysis (Supplemental Fig. S4C). We further dissected the transcriptional response to HS using 150 TPM (Supplemental Fig. S4D) and 200 TPM (Supplemental Fig. S4E) cutoffs, respectively, and found that the impact was only significant in highly expressed genes. Our results suggest that heat stress at the bicellular stage has a large effect mostly on highly expressed genes and therefore critical for sperm cell development.

### Heat stress mis-regulates cell cycle control genes in sperm cells, but does not affect their DNA content

We next explored in more detail which genes are differentially regulated by HS during sperm cell formation. In plants, the SCF E3-ligase complex formed by the S-Phase Kinase-associated Protein1 (SKP1; Zm00001eb404320), cullin CUL3A (Zm00001eb254590), a F-box protein (Zm00001eb187770), F-box protein GID2 (Zm00001eb245180) and Ring-Box Protein 1A (RBP1-Zm00001eb188540) plays a key role integrating developmental end environmental responses (Ban and Estelle, 2021). Notably, all members of the SCF complex showed upregulation of more than 10-fold (Fig. 7A). Moreover, while most members of the anaphase promoting complex/cyclosome APC/C, another multimeric E3-ligase, are not expressed in sperm cell, (Supplemental Table S4), APC10a (Zm00001eb342630) and CCS52A1 (Zm00001eb001710) were mis-regulated after HS (Fig. 7B). SAMBA (Zm00001eb024740) and CCS52B (Zm00001eb139790), other members of the APC/C complex, showed expression in sperm cell, but transcript levels did not change in response to heat stress (Supplemental Table S4). Since the SCF and APC/C complexes regulate cyclins and cyclin-dependent kinases (CDKs) controlling the progression of cell cycle, we searched for maize CDK genes whose expression pattern changed in response to HS. Out of the total number of cyclins and CDKs described in maize only three genes were differentially expressed in maize sperm cells after HS: ZmCycA1 (Zm00001eb227730), ZmCycD4 (Zm00001eb038430) and ZmCDKB2 (Zm00001eb035350) (Fig. 7C). All three genes were induced. Noteworthy, these three genes were also the only ones expressed in maize sperm cells under control conditions. In summary, we found that heat stress induced upregulation of components of the SCF and APC/C E3-ligase complexes, which are known to control the degradation of cell cycle regulators (cyclins and CDKs) to allow G1-to-S transition as well as S-to-G2 transition (Kim et al., 2008; Villajuana-Bonequi et al., 2019) and thus promoting cell cycle progression (Fig. 7D).

**Figure 7.**
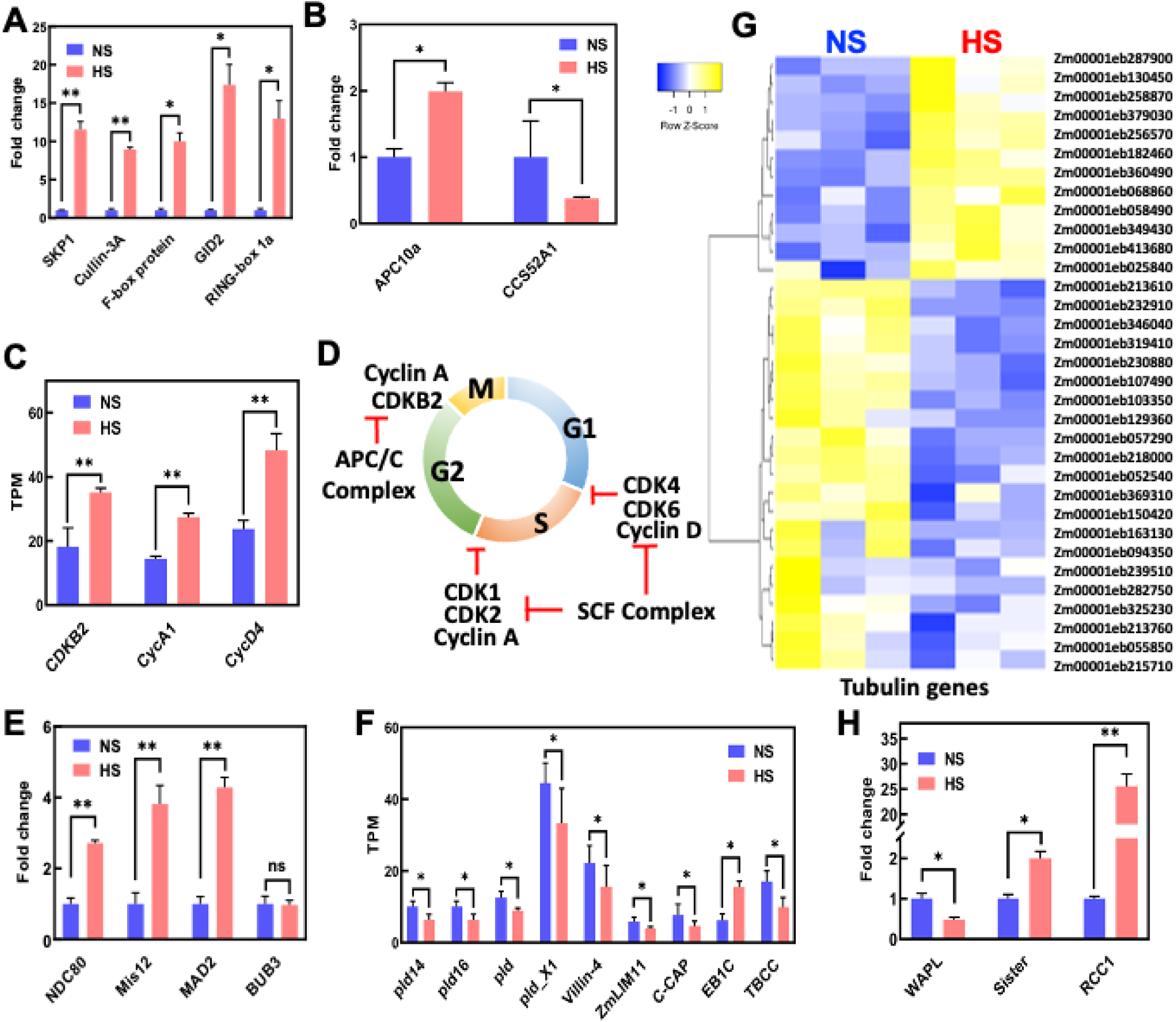
KP1-CUL1-F-Box-Protein (SCF) E3 Ubiquitin Ligase complex and spindle assembly check point (SAC) genes are up-regulated in sperm cells after heat stress was applied during the bicellular stage of pollen development. **(A)** Induction of the SCF members genes. **(B)** Mis-regulation of the APC/C complex genes. **(C)** Up-regulation of cyclins. **(D)** Illustration of the cell cycle during pollen mitosis II (PM II) and its regulation by cyclins and the SCF complex. **(E)** Up-regulation of SAC gene members after heat stress. NDC80, component of the kinetochore complex. **(F)** Down-regulation of microtubule-associated genes. **(G)** Mis-regulation of tubulin-associated genes. **(H)** Mis-regulation of genes involved in mitosis progression during the metaphase to anaphase transition. WAPL, Wings apart-like protein homolog. RCC1, regulator of chromosome condensation 1. Sister, Sister chromatid cohesion 1. One asterisk indicates significant difference at P < 0.01; two asterisks indicate significant difference at P < 0.001; one-tailed *t*-test comparing HS to NS samples. n = 3 biological replicates, each with 3 technical replicates.

We next measured the DNA content of sperm cells to elucidate whether mis-expression of cell cycle regulators may affect DNA replication and ploidy level of sperm cells. In *Arabidopsis*, it has been shown that cold and heat stress disrupts genomic ploidy consistency, generating the formation of diploid and polyploid pollen (De Storme et al., 2012; Lei et al., 2020). Therefore, we performed comparative ploidy level measurements by using flow cytometry on sperm cells isolated from mature pollen previously heat stressed at the bicellular stage. As a control we used nuclei isolated from leaf tissue and observed the 2C and 4C peaks that are typical for this type of tissue indicating diploidy (Supplemental Fig. S5A). Only one dominant 1C peak was observed in NS sperm cells (Supplemental Fig. S4B). Similarly, sperm cells from HS pollen showed one single and sharp 1C peak indicating haploidy and lack of aneuploidy (Supplemental Fig. S5C). These findings demonstrate that sperm cells still contain a normal DNA content, but it does not exclude DNA/chromatin defects that occurred due to improper cell cycle checkpoint controls.

### Heat stress activates spindle assembly check point and meta-to anaphase transition genes in sperm cells

Since our data points toward mis-regulation of genes involved pollen mitosis II, we searched for developmental transitions involved in this process. During the transition from metaphase to anaphase, where chromosomes aligned at the equator of the cell and migrate to the poles, a developmental process ensures that all chromosomes are aligned at the same level. This process is controlled by the spindle assembly checkpoint (SAC) signaling complex, which regulates proper partitioning of chromosomes to daughter cells during mitosis. SAC signaling is a mechanism that it is only active when chromosomes are not properly attached to the kinetochores and thus delays progression of the cell cycle until all kinetochores are correctly assembled (Lara-Gonzalez et al., 2021). We found upregulation of genes encoding Nuclear Division Cycle 80 (Ndc80, Zm00001eb040130), minichromosome instability 12 (Mis12, Zm00001eb433940) and mitotic arrest deficient 2 (Mad2, Zm00001eb425240) in sperm cells generated during HS treatment (Fig. 7E). These genes are part of the SAC signaling pathway. Another member of SAC signaling, budding uninhibited by benzimidazoles 3 (BUB3, Zm00001eb151660), did not change in response to HS (Fig. 7E). In conclusion, these results indicate that HS activates SAC signaling and possibly also the generation and regulation of the spindle apparatus. Therefore, we searched for microtubule-related genes in our sperm cell data sets. We identified nine microtubule-related genes with TPM vales higher than 5 and classified them in two groups, (i) microtube-related signaling and microtubule-functions (Fig. 7F). Four of the microtube-related signaling genes belong to the Phospholipase D family. Phospholipase D activity has been correlated with many physiological processes including organization of the cytoskeleton, endocytosis and exocytosis, but also stress responses during development and immunity (Ali et al., 2022). All four Phospholipase D genes were downregulated in response to heat stress (Fig. 7F). Similarly, VILLIN3 (Zm00001eb389350), a gene required for the generation of actin filament bundles (Van Der Honing et al., 2012) was downregulated after heat stress at the bicellular stage of pollen development. Another microtube-related signaling gene encodes a LIM (Lineage-11 (LIN-11), Insulin-1 (ISL-1), and Mechanotransduction-3 (MEC-3) - Zm00001eb268250) gene. LIMs are cytoskeleton-associated proteins that inhibit actin filament depolymerization and cross-links filaments in bundles (Thomas et al., 2007). Downregulation of this gene was also found after heat stress (Fig. 7F). Another gene in the microtube-related signaling category encodes an adenylyl cyclase-associated protein (C-CAP, Zm00001eb040800), which similarly as all the others was downregulated in response to heat stress (Fig. 6F). The only two differentially expressed genes involved directly in microtubule function showed opposite transcriptional expression: while microtubule-associated protein EB1C (Zm00001eb044540) was upregulated, tubulin binding cofactor C (TBCC, Zm00001eb236410) was downregulated in HS sperm cells (Fig. 7F). We further explored a full set of structural tubulin genes differentially expressed after heat stress and found that more than 63% of them were downregulated (Fig. 7G).

Another set of genes important for the transition from metaphase to anaphase are Wings Apart-like Protein homolog (WAPL, Zm00001eb220920), Regulator of Chromosome Condensation 1 (RCC1; Zm00001eb049950) and Sister chromatid cohesion 1 (Sister; Zm00001eb412050) (Fig. 7H). WALP induces cohesin dissociation from DNA allowing the progression of the mitotic cell cycle, which is modulated by Sister (Crawley et al., 2016). We found that WALP was downregulated after heat stress, while Sister is upregulated (Fig. 7H). RCC1, another important gene whose product associates to chromatin dynamically controlling metaphase-to-anaphase transition was highly upregulated after heat stress (Fig. 7H). Collectively, our results show that heat stress impact sperm cell development by targeting spindle assembly check point and metaphase-to-anaphase transition genes whose mis-regulation might cause defective sperm cells.

## DISCUSSION

Heat stress (HS) directly affects plant reproduction processes by decreasing the viability of reproductive cells ultimately leading to fertilization failures and sterility, and thus declining plant productivity (Chen et al., 2016; Begcy et al., 2019; Chaturvedi et al., 2021; Kumar et al., 2023). Here, we show that HS applied exclusively either at the unicellular or bicellular stages of pollen development results in a strong reduction in seed set and yield. Reduction in seed set upon exposure to high temperatures during the plant reproduction phase have been observed in many field crops (Prasad et al., 2017; Sun et al., 2018; Begcy et al., 2019; Wang et al., 2020), but the underlying molecular mechanisms affected are largely unclear. Defects in pollen viability and tube growth are common alterations after HS leading to decrease in productivity (Endo et al., 2009; Begcy et al., 2019). However, our results showed that imposing HS during the bicellular stage of pollen development on maize did not result in noticeable changes in morphology, vitality, and germination capabilities.

Formation of the two sperm cells is the central developmental process occurring at the bicellular stage. Disruption of spermatogenesis in mammals due to HS has been shown to negatively impact male reproduction altering sperm morphology and leading to DNA fragmentation (Hansen, 2009; Garcia-Oliveros et al., 2020; Capela et al., 2022). In plants little is known on the impact of high temperatures for sperm cell formation. In *Arabidopsis*, sperm-less pollen tubes were obtained after knocking out two bHLH transcription factors, *drop1* and *drop2* (Zhang et al., 2017), indicating that proper pollen tubes can be formed lacking sperm cells, but probably also containing defective and non-functional sperm cells. Pollen tubes in maize are also formed after HS application at the bicellular stage, but sperm cells were not transported. We have shown that tubulin and other cytoskeletal protein genes are mis-regulated and protein levels reduced after heat stress. Notably, a decline in the number of functional proteins, particularly those involved in cytoskeletal organization was previously also shown in mature HS tomato pollen (Keller et al., 2018). While transport of sperm cells and vesicles in pollen tubes is largely actin-dependent, microtubules and their kinesin motors play among others an important role of linking the sperm cells with the vegetative nucleus forming the male germ unit (McCue et al., 2011; Cai, 2022). Their mis-expression can well explain the observed transport defect.

Among the molecular mechanisms that ensures proper sperm cell development is the SAC (spindle assembly checkpoint) complex. The SAC is a surveillance mechanism that ensures error-free chromosome segregation by blocking the metaphase-to-anaphase transition during mitosis and meiosis. It is only active when chromosomes are not correctly attached to spindle microtubules emanating from opposing poles (Komaki and Schnittger, 2016; Komaki and Schnittger, 2017; Lara-Gonzalez et al., 2021). We found that HS activates SAC signaling in sperm cells. Since the SAC is likely also necessary to ensure genome stability during sperm cell formation (PM II), it is not unlikely that its mis-regulation may lead to chromosome defects. Our flow cytometry data indicated that these cannot be dramatic as the ploidy level of sperm cells remained unchanged and aneuploidy could not be detected using this method. This could also be because of a phenomenon called mitotic slippage or SAC adaptation: during mitotic slippage, the control of the SAC to delay mitosis disappears if the time taken to form proper chromosomal attachments to spindle microtubules is too prolonged. Thus, the mitotic process continues, and the cells will eventually divide (Sinha et al., 2019). Since mature pollen grains after HS application at the bicellular stage generate functional tubes and contain two sperm cells, but seed formation is highly reduced, it is not unlikely that defective sperm cells are the cause due to genome/chromosome defects that occurred during HS at PM II. Genomic sequencing of single sperm cells could now elucidate whether the sperm cells contain defective DNA and/or chromosomes.

Other important components for functional sperm cell development are the cytoskeleton and microtubule organization. In *Arabidopsis*, high temperature conditions were shown to interfere with the configuration of α-tubulin, affecting the construction of the spindle and phragmoplast during male meiosis I and II (Lei et al., 2020). Similarly, in tobacco cells, HS affects the microtubules of the mitotic spindle and phragmoplast, resulting in split spindles, altered microtubule asters, and elongation of the phragmoplast (Smertenko et al., 1997). Notably, when microfilaments essential for sperm nuclear migration in the egg cell cytoplasm is disrupted in *Arabidopsis*, the sperm cell nucleus failed to fuse with either the central and egg cell nucleus, respectively (Kawashima et al., 2014). In CENH3 *Arabidopsis* mutants, high-temperature induced pollen reprogramming development to an embryogenic one by reorganization of microtubules (Ahmadli et al., 2023). This suggests that as the temperature increases, the stability of microtubules and chromosomes decreases, as evidenced also in this study by the reduced signal intensity of the centromeric histone CENH3 marker line under heat stress. Of course, also other genes like those required for proper transcription and translation were mis-regulated after HS explaining reduced protein levels and supporting the hypothesis that non-functional sperm cells are formed. This has to be investigated in future experimentation, which is still a very challenging task in species containing thick female tissues like maize.

In summary, we show that HS applied at the bicellular stage of pollen development leads to mis-regulation especially of cell cycle regulatory genes and likely impairs also progression during the metaphase to anaphase transition by the hyperactivation of SAC, which altogether probably causes defective sperm cells and their male germ units that cannot be properly transported in the pollen tube and do not arrive at the female gametophyte to execute double fertilization. Whether and how mis-expression of the described genes is avoided in HS tolerant plants will be an exciting task for the future.

## MATERIALS AND METHODS

### Plant material, reporter lines and growth conditions

In this study, maize (*Zea mays*) inbred line B73 and sperm cell-specific α-tubulin-YFP (Kliwer, et al., 2009) as well as mRuby3-CENH3 marker lines were used for heat stress studies. The centromeric mRuby3-CENH3 (GEX3p:mRuby3-CENH3) line was generated as follows: 1.3 kbp upstream of the open reading frame (ORF) of the sperm cell-specific *ZmGEX3* gene (Chen et al., 2017; gene ID Zm00001eb161540) was cloned together with the ORF of a mRuby3-CENH3 fusion protein gene into the vector pTF101.1 provided by the maize transformation platform at Iowa State University. Transgenic maize plants were generated using hybrid embryos of inbred lines Hi-IIA and Hi-IIB. All maize seeds were germinated in an incubator, transferred after 10 days to a greenhouse into larger pots (10 cm diameter, 10 seedlings per pot) containing a standard substrate and soil mixture (1:1, v/v). After three weeks maize seedlings were planted into 10 L pots grown under controlled conditions of 16 h of light at 26°C ± 2°C and 8 h of darkness at 21°C ± 2°C, and a constant air humidity of 60% to 65%. Supplemented light of ∼20,000 lux was provided to adjust day length duration. An automated temperature-water-based irrigation system was used to supply water according to plant consumption in a time-based preprogrammed schedule. Plants were fertilized twice a week with 2% (w/v) Hakaphos and monitored throughout their vegetative and reproductive development. Marker line plants were genotyped before transfer into larger pots. All plants were monitored throughout their entire vegetative and reproductive development.

### Heat stress treatment at the unicellular and bicellular stages

Heat stress (HS) was applied at the uni- and bicellular stages of maize pollen development. Identification of pollen developmental stages was performed using the Leaf Collar Method as described previously (Begcy and Dresselhaus, 2017). After reaching either of the stages, maize plants were transferred to walk-in growth chambers. For heat stress, growth chamber day/night temperature conditions were set at 35° for 16h light and 25°C darkness with 60% air humidity at 25,000 lux for 48 h. Correspondingly, non-stressed (NS) plants were maintained at a 25°/21°C day/night temperature regime and 60% humidity at 25,000 lux in a control chamber. After 48 h exposure to heat stress, all plants were maintained under control conditions as described above until pollen maturation. Pollen was either collected directly afterwards for morphological and physiological analyses or at pollen maturity for germination and seed set assays. Samples for biochemical and RNA-Seq analysis were also collected from pollen at maturity.

### Pollen germination assays

Maize plants used for pollen germination experiments were placed close together to maintain similar conditions and to ensure high quality of pollen harvested. For *in vitro* pollen germination, solid medium was prepared by mixing 2x Pollen Germination Medium (PGM: 20% sucrose; 0.005% H_3_BO_3_; 20mM CaCl_2_; 0.1 mM KH_2_PO_4_; 12% PEG4000, pH=5) and an equal volume of autoclaved 1.2% NuSieveTM GTGTM agarose (Lonza) to a final agarose concentration of 0.6%. A total of 3 mL of PGM medium mixture was pipetted into a 35 mm petri dish and gently agitated horizontally to obtain a thin and evenly distributed medium layer after 10 min solidification at room temperature (RT). Freshly collected pollen from NS and HS plants was obtained by softly shaking new released florets into petri dishes containing solid PGM. Pollen was germinated at room temperature (22-23°C) in a wet and dark chamber. The pollen germination status was monitored after 40-45 min on PGM and visualized using a Nikon Eclipse 1500 microscope with a 4x objective (Plan Fluor DL 4x/0.13, PHL) equipped with a Zeiss AxioCam MRM monochromatic camera.

*In vivo* pollen germination was carried out as previously described (Begcy et al., 2019) with some modifications. Before silking, ears from NS maize plants were covered using small paper bags to prevent pollen contamination. Fresh pollen grains from NS and HS plants were harvested using a paper bag and pollinated on newly emerged silks. After one hour of *in vivo* pollen germination, 5 cm from the top portion of pollinated silks were cut off and fixed in 9:1 v/v ethanol: acetic acid at 4°C overnight. Fixed samples were rehydrated by a water series. Then, silks were treated with 8 M sodium hydroxide for 2-4 h to clear and soften the tissue. Softened silks were washed 2-4 times using water. Staining was carried out using aniline blue staining solution (0.1% aniline blue; 0.1 M K_2_HPO_4_·3H_2_O, pH=11) overnight at 4°C. Samples were washed and mounted using fresh staining solution on a slide with a cover slip and analyzed under fluorescence microscope ZEISS Axio Imager 2 with a 20x objective (Plan-Apochromat 20x/0.8 M27) at 350∼400nm (UV) excitation.

### Maize sperm cell isolation

A discontinuous percoll density gradient centrifugation method was used as previously described (Dupuis et al., 1987; Chen et al., 2017) with some modifications. Fresh pollen grains from NS and HS B73 maize plants were harvested and placed into glass petri dishes containing a moist filter paper in the internal part of the lid. Pollen was allowed to pre-hydrate at room temperature for at least 2 h. Then, pollen was immersed in 550 mOsmol·kg^−1^ H_2_O mannitol solution (100 mg pollen/mL solution) and incubated on a platform shaker with slow agitation (80 rpm) for 1-2 h. Pollen lysates were filtered in a 50 mL Falcon conical tube equipped with a pluriStrainer 30 µM polyester mesh (PluriSelect) and a connector ring (PluriSelect) allowing force filtering of samples using manual pressure with a syringe. Subsequently, a cleared lysate obtained from the filtering step was layered on top of a discontinuous 3-phase percoll gradient consisting of 30%/20%/15% (v/v) percoll in 550 mOsmol·kg^−1^ H_2_O mannitol solution. Centrifugation was performed at 12,000×g for 1 h at 4°C. After density gradient centrifugation, the 20%/30% interphase, where sperm cells accumulated as a faint yellowish line, was collected using a Pasteur pipette and transferred to a 10× volumes 550 mOsmol·kg^−1^ H_2_O mannitol solution, followed by another centrifugation at 2,500×g to wash out pollen organelles and other cytoplasmic contaminants. Pelleted sperm cells were resuspended in 20 μL of 550 mOsmol·kg^−1^ H_2_O mannitol solution and the number of isolated sperm cells was assessed using a cell counting chamber (Marienfeld). Isolated sperm cells were used immediately or shock-frozen in liquid nitrogen and stored at −80°C.

### RNA isolation and RT-qPCR

To ensure high RNA quality, all tubes and tips used during pollen RNA extractions were RNase-free and metal beads were autoclaved. Working bench and pipettes were cleaned with RNase Zap RNase Decontamination Solution (Thermo Fisher Scientific). Total RNA was isolated from NS and HS sperm cells or pollen grains using the Invitrogen™ TRIzol™ Plus RNA Purification Kit (Thermo Fisher Scientific). Approximately 0.2 g pollen were each collected in a 2 mL microcentrifuge tube with one metal balls inside. Samples were frozen in liquid nitrogen and ground to powder using a TissueLyser II (Qiagen). 1 mL of TRIzol™ reagent was added to each sample and vortexed adequately. After 5 min incubation at RT, 200 mL of chloroform was added to each sample. After inverting several times and incubation for 2-3 min, samples were centrifuged for 15 min at 12,000xg at 4°C. Around 600 µL upper colorless layer supernatant containing RNA was transferred to a new tube and mixed with equal volume of 70 % ethanol by inversion. The mixture was transferred into spin cartridges provided by the kit and centrifuged at 12,000 g for 15 seconds. To remove DNA contamination, RNase free DNase solution (Qiagen) were added to each column and centrifuged. RNA on the membrane were washed by sequentially washing buffers supplied with the kit. Total RNA was eluted out from membranes by adding 50 µL pre-warmed 60°C RNase-free water for obtaining high RNA yield. For quantification and quality control, isolated RNA was photometrically analyzed using the Nanodrop ND 100 system (Thermo Fisher Scientific) and stored at -80°C. Complementary DNA (cDNA) synthesis was performed using 1 μg of total RNA using Oligo (dT) 18 primer and Reverse Transcriptase RevertAid™ (Thermo Fisher Scientific) according to instruction as previously described (Preciado et al., 2022). For normalization, an ubiquitin gene (gene ID Zm00001eb066940) was used as a control. PCRs were performed using the Master Cycler realPlex2 (Eppendorf) in a 96-well reaction plate according to the manufacturer’s recommendations. Primers are listed in Supplemental Table S6. Cycling parameters consisted of 5 min at 95°C followed by 40 cycles of 95°C for 15 s, 60°C for 30 s, and 70°C for 30 s. Reactions were performed in triplicate for each RNA sample using at least three biological replicates. Specificity of amplifications was verified by melting curve analyses. Results from the Master Cycler realPlex2 detection system were further analyzed using Microsoft Excel (Kim et al., 2021). Relative amounts of mRNA were calculated from threshold points (Ct values) located in the log-linear range of RT-qPCR amplification plots using the 2-^ΔΔCT^ method (Livak and Schmittgen, 2001).

### DAPI microscopy

Isolated pollen grain at different stages using the Leaf Collar Method (Begcy and Dresselhaus, 2017) were placed on glass slides containing 1 mg/mL DAPI (4′,6-diamidino-2-phenylindole) solution in 1× phosphate-buffered saline (PBS; 0.8% [w/v] NaCl, 0.002% [w/v] KCl, 0.014% [w/v] Na_2_HPO_4_, and 0.0024% [w/v] KH_2_PO_4_) and observed using a Zeiss Axio Imager Z1 microscope equipped for structured illumination (Apotome2) and with a Zeiss AxioCam MRM monochromatic camera. DAPI fluorescence was observed using a filter set with a wavelength of 359 nm.

### Pollen vitality assays

Freshly harvested mature pollen from NS and HS plants were subjected to cytological analysis to assess their metabolic activity. Pollen grains were stained with fluorescein diacetate (FDA) as previously described (Begcy et al., 2019) and with Alexander staining then visualized using a Zeiss Axio Imager Z1 microscope (Apotome) equipped with a fluorescein isothiocyanate (FITC) filter, which allows the detection of fluorescein. Excitation and emission filters were set at 450 nm and 520 nm wavelength, respectively (Zeiss). Pollen grains that exhibited fluorescence in the FITC channel were considered metabolically active.

### Library preparation and Illumina RNA sequencing

Library preparation and RNA sequencing were conducted following the Illumina TruSeq Stranded mRNA Sample Preparation Guide, the Illumina HiSeq 1000 System User Guide (Illumina), and the KAPA Library Quantification Kit-Illumina/ABI Prism User Guide (Roche). In brief, 250 ng of total RNA of NS and HS sperm cells were used for purifying polyA^+^ containing mRNA molecules using poly-T oligo-attached magnetic beads. Three biological replicates were used per condition. After purification, mRNA was fragmented to achieve an average insert size of 200–400 bases using divalent cations under elevated temperature. Resulting cleaved RNA fragments were reverse transcribed into first-strand cDNA using reverse transcriptase, Actinomycin D and random hexamer primers. Subsequently, blunt-ended second-strand cDNA was obtained using DNA Polymerase I, RNase H, dUTP, and 2′-Deoxyuridine, 5′-Triphosphate (dUTP) nucleotides. Adenylation of the resulting cDNA fragments at their 3′ ends were performed followed by ligation with indexing adapters. PCR enrichment was used to subsequently create specific cDNA libraries. The KAPA SYBR FAST ABI Prism Library Quantification Kit (Roche) was used to quantify the libraries. Equimolar amounts of each library were used for cluster generation on the cBot using the Illumina TruSeq PE Cluster Kit v3. Sequencing was carried out on a HiSeq 1000 instrument using the indexed, 50 cycles paired-end read (PR) protocol and the TruSeq SBS v3 reagents according to the Illumina HiSeq 1000 System User Guide. Bcl files obtained from image analyses and base calling were converted into fastq files using the bcl2fastq v2.18 software (Illumina). Library construction and RNA-Seq were performed at the Regensburg University service facility KFB (Competence Center for Fluorescent Bioanalytics).

### Data processing, mapping, differential expression, and statistical analysis

RNA sequencing paired-end reads quality was assessed using FastQC, followed by trimming and filtering using Trimmomatic v.0.35 (Bolger et al., 2014). Processed paired-end reads were mapped and aligned using the STAR v. 2.5.2 program (Dobin et al., 2013) and the maize reference genome sequence AGPv3 assembly using annotation release-5b+ (corresponding to Gramene AGPv3.27). Alignment results were summarized per gene by identifying read pairs aligned to exonic regions with featureCounts (Liao et al., 2014). Reads overlapping with more than one feature (i.e. gene region) were excluded from the analyses. We performed variance stabilizing transformation of raw counts and analyzed non-merged technical replicates using R package DESeq2, to evaluate the presence/absence of batch effect and outliers (Love et al., 2014). Differential expression analysis between NS and HS pollen samples was conducted using fold change >1.5 and P <0.05 (after the false discovery rate adjustment for multiple testing) for the null hypothesis as threshold using DEseq2 (Love et al., 2014).

To elucidate transcriptional correlations, the cor-function was implemented in the R package WGCNA (Langfelder and Horvath, 2008) to identify genes with shared expression profile. Genes exhibiting similar expression profiles were considered as seed candidates to obtain direct and indirect interactions using the STRING v11 database (Szklarczyk et al., 2019). A threshold a high confidence score of 0.7 was used to construct cluster correlation analysis. Only interactions with high levels of confidence were extracted from the database and considered as valid.

### Flow cytometry analysis

Sperm cells were harvested and used for isolation as described above. Isolated sperm cells were incubated in staining solution (CyStain UV Ploidy, Sysmex) for 5-15 min in the dark (Grossman et al., 2021). After staining, samples were further purified using 50 μm filters and their DNA content was analyzed via flow cytometry using the CyFlow Space system (Sysmex).

### AP-MS analysis of proteins

Mature pollen grains were used for proteomic analyses. For protein extraction, 50 mg of pollen grains were ground with liquid nitrogen using a mortar and dissolved in 250 μL of ice-cold extraction buffer (50 mM Tris/HCl, 150 mM NaCl, 0.1% sodium deoxycholate, 0.1% Triton-X100, 1 mM PMSF, pH 8.0). Proteomic analyses were performed as described (Antosz et al., 2020). In brief, total proteins from pollen samples were separated on a 10% SDS-PAGE gel containing the PageRuler™ Prestained Protein Ladder as a molecular weight marker (Thermo Fisher Scientific). After in-gel digestion with trypsin (Promega), the resulting peptide mixtures were loaded onto the UltiMate 3000 RSLCnano system, which was connected to an Orbitrap Elite hybrid mass spectrometer (MS, Thermo Fisher Scientific). MS data were acquired using a data-dependent strategy that selected the top 10 precursors for higher-energy collisional dissociation (HCD) fragmentation. The analysis of MS raw data files was performed using the Proteome Discoverer software (Thermo Fisher Scientific; version 1.4), utilizing in-house Mascot (Matrix Science; version 2.6) and Sequest search engines. MS/MS ion searches were conducted against the UniProt (The UniProt Consortium et al., 2023) and the Gramene (Tello-Ruiz et al., 2022) protein databases for maize (*Zea mays L*). Post-processing of search results was performed using Percolator (Spivak et al., 2009). Peptides with a q value < 0.01, rank 1, and a minimum length of six amino acids were only considered. Protein abundance was determined using the protein area calculated by the Proteome Discoverer software (Thermo Fisher Scientific; version 1.4).

### Statistical analysis

Statistical analyses were performed using R software/environment. For all measurements, data from at least four independent experiments were used. Data represented mean and median values including standard deviations. Wilcoxon’s singed-rank test was used to compare gene expression between HS and NS plants (Kim et al., 2021). Differences in means were considered significant at p-value < 0.05.

## Author contributions

K.B. and T.D. conceived the experiments. X.L. performed the experiments. A.B. supervised the proteomic analysis. K.B, T.D. and X.L. analyzed the data. K.B. wrote the manuscript with input from all authors. All authors read and approved the manuscript.

## One sentence summary

Pollen exposed to moderate heat stress for two days at the bicellular stage generate defective sperm cells containing mis-regulated replication and other cell cycle-associated as well as cytoskeletal genes ultimately affecting their transport inside pollen tubes.

## Acknowledgments

We would like to thank Dr. Kamila Kalinowska for generating the CENH3 marker construct and acknowledge Armin Hildebrand for plant care.

## SUPPLEMENTAL DATA

**Supplemental Figure S1.** Heat stress at the unicellular stage of maize pollen development decreases pollen viability.

**Supplemental Figure S2.** Heat stress during the bicellular stage of pollen development in maize does not affect pollen morphology and viability.

**Supplemental Figure S3.** Gene ontology enrichment analysis of genes differentially expressed in maize pollen in response to heat stress.

**Supplemental Figure S4.** Heat stress alters transcriptional gene expression of highly expressed genes.

**Supplemental Figure S5.** Heat stress at the bicellular stages does not affect DNA content of sperm cells in maize.

**Supplemental Table S1.** List of identified proteins involved in cytoskeletal organization.

**Supplemental Table S2.** Summary of sperm cells samples including number of cells per replicate, aligned pairs mapped unique and percentage of reads per trimmed reads. Alignment to the AGPv3 assembly using annotation release-5b+ (corresponding to Gramene AGPv3.27).

**Supplemental Table S3.** Differentially expressed genes between non-stressed and heat stressed samples during the tetrad stage of pollen development.

**Supplemental Table S4.** List of genes used for gene cluster correlation analysis.

**Supplemental Table S5.** TPM vales of APC/C members under nonstressed and heat-stressed conditions.

**Supplemental Table S6.** List of primers used in study.

